# SENSV: Detecting Structural Variations with Precise Breakpoints using Low-Depth WGS Data from a Single Oxford Nanopore MinION Flowcell

**DOI:** 10.1101/2021.04.20.440583

**Authors:** Henry CM Leung, Huijing Yu, Yifan Zhang, Wing Sze Leung, Ivan FM Lo, Ho Ming Luk, Wai-Chun Law, Ka Kui Ma, Chak Lim Wong, Yat Sing Wong, Ruibang Luo, Tak-Wah Lam

## Abstract

Structural variation (SV) is a major cause of genetic disorders. In this paper, we show that low-depth (specifically, 4x) whole-genome sequencing using a single Oxford Nanopore MinION flow cell suffices to support sensitive detection of SV, in particular, pathogenic SV for supporting clinical diagnosis. Existing SV calling software, when using 4x ONT WGS data, often fails to detect pathogenic SV especially in the form of long deletion, terminal deletion, duplication, and unbalanced translocation. Our new SV calling software SENSV is able to achieve high sensitivity for all types of SV and a breakpoint precision typically ±100 bp, both features are important for clinical concerns. The improvement achieved by SENSV stems from several new algorithms. We evaluated SENSV and other software using both real and simulated data. The former was based on 24 patient samples, each diagnosed with a genetic disorder. SENSV found the pathogenic SV in 22 out of 24 cases (all heterozygous, size from hundreds of Kbp to a few Mbp), reporting breakpoints within 100 bp of the true answers. No existing software can detect the pathogenic SV in more than 10 out of 24 cases, even when the breakpoint requirement is relaxed to ±2,000 bp.

## Introduction

Structural variation (SV) refers to the changing of copy number, orientation or position of DNA segment; it occurs in the form of deletion, duplication, inversion, unbalanced/balanced translocation. Many genetic disorders are caused by SV, which is often heterozygous in nature. Accurate detection of SV is important in clinical diagnosis. In this paper, we consider SV with size of at least one thousand base pairs^1^ and in particular pay attention to pathogenic SV which often involves over a hundred thousand base pairs.

Latest SV detection software tools^2–7^ exploit long (yet highly-erroneous) reads from the third-generation sequencing (3GS) technologies to achieve higher sensitivity than the clinically used methods (including CGH array and low-pass NGS approach) or even the high-depth NGS approach^8–10^; furthermore, the breakpoint precision can be enhanced from ±50 Kbp^11,12^ to ±2,000 bp or even ±100 bp in some cases. Sniffles^2^ and SVIM^4^ are the representative software tools here. They were designed to detect SV using WGS data with high depth (i.e., 30x), which currently costs a few thousand US dollars using Oxford Nanopore Technologies (ONT), or even higher using Pacbio Technologies^13^. This cost is too high compared with existing methods used in clinical settings. At present, ONT can support 4x WGS with a single MinION flowcell (or a fraction of a PromethION flowcell), costing a few hundred US dollars. NanoVar^14^ was the first to demonstrate success in using low-depth ONT data, it recommends using at least 4x ONT data for homozygous SV and at least 8x for heterozygous SV.

Using 4x ONT data, existing software can still detect an acceptable number of SVs when benchmarked against a normal reference sample like HG002, but they often fail to detect pathogenic SVs when evaluated using patient samples. Notice that almost all the confirmed SVs in HG002 are deletions with less than 100 Kbp, yet known pathogenic SVs are much longer, often more than 100 Kbp (over 95% of confirmed pathogenic SVs in dbVar have more than 100 Kbp in length). Existing software can detect deletion but the sensitivity drops drastically when the size exceeds 100 Kbp; and they have difficulty in detecting duplication, terminal deletion^15^ and unbalanced translocation.

To overcome the difficulty in detecting pathogenic SV using 4x ONT data, we devised a new software tool called SENSV, which can detect heterozygous SV with better sensitivity for all sizes (including over 100Kbp) and for all types of SV. Furthermore, SENSV achieves a breakpoint precision typically ±100 bp. When diagnosing genetic disorders, it is useful to obtain more precise breakpoints^16^ so as to determine the exact genes affected by an SV and its functional impact. Ideally, if breakpoint precision can be improved to ±100 bp, the target pathogenic SV in other samples from the patient or the patient’s relatives can be verified using PCR at very low cost.

The improvement achieved by SENSV stems from three new algorithms for recovering misaligned split reads, boosting the confidence of true alignment and refining breakpoints. We evaluated SENSV together with NanoVar, Sniffles, and SVIM using the data from 24 patients diagnosed with genetic disorder (DNA samples were obtained from the Clinical Genetic Service of the Department of Health, Hong Kong). In each case, the pathogenic SV was known to be a heterozygous SV and had been clinically confirmed. Each patient sample was sequenced using a single MinION flowcell (average depth 4.2x). Among the 24 cases, 16 of them are difficult cases, where the pathogenic SV in the form of long deletion (i.e., over 100 Kbp), duplication, terminal deletion and unbalanced translocation, and the remaining 8 cases involve an inversion or balanced translocation which are relatively easy to detect (as the expected number of reads covering the boundary of these two types of SVs is double than that of other SVs^17^). SENSV outperformed the others in detecting pathogenic SV; it detected 14 out of the 16 difficult cases (22 out of 24 in total), all with breakpoint precision within 100 bp. Other software detected at most 3 difficult cases and 9 cases in total (two more difficult cases can be detected if the breakpoint precision is relaxed to ±2,000 bp in accordance with the GIAB evaluation practice^18^). SENSV’s improvement stems primarily from its ability to recover misaligned split reads.

Besides patient samples, we also sequenced the normal reference sample HG002^18^ using a single MinION flowcell (depth 3.7x) and benchmarked SENSV and other software. HG002 contains 532 SVs confirmed by GIAB (with size at least 1 Kbp). SENSV detected 390 (73% of 532) GIAB-confirmed SVs with breakpoint precision ±100 bp; the total number of SVs predicted is 3,644. SVIM has the second best performance; it predicts more SVs (6,650 in total) but detects slightly fewer GIAB-confirmed SVs (379; 71% of 532). When breakpoint precision was evaluated using GIAB’s practice of ±2,000 bp, SENSV and SVIM could detect 432 (81% of 532) and 428 (80% of 532) GIAB-confirmed SVs, respectively. Notice that all the 532 confirmed SVs, except one, are short deletion with size less than 100 Kbp.

As revealed by the patient data, pathogenic SVs in the form of long deletion (over 100 Kbp), duplication, terminal deletion or unbalanced translocation are relatively difficult to detect. Yet such SVs are not found in HG002 (except one long deletion). Thus we further benchmarked the software for these types of SVs using simulated data. We created two simulated 4x ONT datasets based on HG19, each implanted 35 SVs of 7 types: short deletions, long deletions, duplications, terminal deletion, unbalanced translocation, inversion and balanced translocation. All these SVs were heterozygous and non-overlapping, either randomly selected from dbVar or generated using RSVSim (when dbVar does not contain enough known SVs of these types).

SENSV outperformed other software in detecting the 70 implanted SVs in the two datasets. When the breakpoint resolution was required to be ±100 bp, SENSV detected 58 out of the 70 SVs (31 out of 40 difficult cases) and predicted a total of 255 SVs. The other three tools detected 33, 33 and 32 SVs (12, 10 and 14 difficult cases and predicted 494, 455 and 1,140 SVs); note that terminal deletions and unbalanced translocations are often missed by existing software tools. More details are given in Result and Discussion.

Intuitively, an SV will give rise to a split-read alignment at its breakpoint in the sense that i) a read is aligned with a big gap, or ii) the prefix and suffix of the read aligned separately to different parts of the genome. For the former case, existing alignment software often relies on reducing the gap penalty in aligning a read to favor finding a big gap^2,19^. However, a split-read alignment might still be missed if the breakpoint occurs towards the end of a read, or if the SV is too large (say, hundreds of Kbp for many pathogenic SVs) for the gap penalty to be worth considering. For the latter case, in order to align the prefix and suffix of a read separately, it requires the read to have a sufficient long prefix and suffix on each side of the SV. Using low-depth data, there may not be any such reads. In summary, it is difficult to obtain a confident split-read alignment using low-depth WGS especially for unbalanced SV which have fewer split-reads than balanced SV.

On the other hand, as the error rate of 3GS data is high (10+% for ONT), a false positive split-read alignment may be introduced. When using low-depth data, there might not be sufficient correctly aligned reads in the same region to confirm a false positive, thus causing large number of predicted SVs.

To overcome the challenges, SENSV introduces an extra step to analyze the sequencing depth of each 10K genome interval against the average sequencing depth of 24 references when sequenced using ONT for 4x WGS (noted that we precompute the average and standard deviation of the 24 references in advance). Intervals with unexpectedly high or low sequencing depth might cover a possible SV. The related reads are then realigned using an SV-aware aligner, called SV-DP, which allows zero penalty for one gap and can recover most of the poorly aligned or misaligned split-read alignments. SV-DP is based on highly optimized dynamic programming to circumvent the efficiency bottleneck in finding a big gap. The sequencing depth analysis and SV-DP contributed to the success of SENSV in detecting the SV from the real ONT data of the 24 patients.

To improve the precision of detected SVs, we relied on three strategies. First, for a false-positive unbalanced SV caused by misalignment, the related interval is likely to have a sequencing depth consistent with the 24 references. This gives SENSV an effective filter. Second, we evaluated each detected SV by realigning the related reads to an altered reference sequence with the conjectured SV inserted into the reference genome. The realignment should show a significant improvement compared to the alignment with the original reference; otherwise, it is likely a false positive. Third, we implemented a filter using metrics including alignment score, sequencing depth and allele frequency of the detected SV to reduce the number of false positive variants.

In addition to having higher sensitivity and precision, SENSV can detect SV with more precise breakpoints (±100 bp) by refining each reported breakpoint using SV-DP and resolving multiple split-reads (if any) with similar breakpoints using realignment.

The details of the above-mentioned algorithms are given in the Method section.

## Results and Discussion

We benchmarked the performance of SENSV and existing software NanoVar^14^, Sniffles^2^ and SVIM^4^ in detecting heterozygous SVs from low-depth ONT WGS data. We used 1) real data from 24 patients with genetic disorders and experimentally verified SVs; 2) HG002 sequenced by a single MinION flow cell; evaluation is based on a recently published SV set; and 3) two simulated datasets each with 35 planted SVs of various types.

### Real data from patients with genetic disorders

The DNA samples of the 24 patients were sequenced using a single MinION flowcell and base-called using the Guppy flip-flop model. The average depth was 4.2x (lowest 2.9x; highest 6.5x). The pathogenic SVs of the 24 samples and their rough positions were diagnosed by medical doctors using conventional methods, including array CGH and karyotype analysis. These SVs were all heterozygous. The first 10 samples involved a simple deletion or duplication occurring at a normal region of a chromosome; the next 6 samples were more complicated, involving an unbalanced translocation or terminal deletion occurring near a highly repeated telomere region. Sample 17 to 24 involved an inversion or a balanced translocation. The (true) breakpoints of SVs were determined based on a detailed analysis by a bioinformatician and a medical doctor; ten samples were selected randomly and the true breakpoints were all confirmed by PCR and/or Sanger sequencing. These true breakpoints were used to evaluate the software tools using SURVIVOR^20^.

The results of the evaluation are shown in Table 1. Samples 1 to 10 (in increasing order of SV length) are each concerned with a simple deletion or duplication. SENSV detected 9 out of 10 SVs, reporting breakpoints within 100 bp of the true breakpoints. The best of the other software detected at most five SVs when we relaxed the breakpoint resolution to 2,000 bp (marked as Y-). Note that the existing software was not designed for detecting heterozygous SV using WGS data with depth as low as 4x, and they might not find sufficient supporting split-read alignments to detect the SV. Table 1 also suggests that the existing software is not sensitive for detecting large unbalanced SVs (length 638 Kbp or more). SENSV can detect large unbalanced SVs using sequencing depth information and recover the precise breakpoint of SV by SV-DP and realignment (samples 4 to 10).

**Table 1.** Comparison of SENSV, NanoVar, Sniffles and SVIM on the ability to detect the pathogenic SV from the 24 patients’ ONT WGS data. Below, “Y” [and “Y-”] mean that a method can detect the pathogenic SV with correct SV type and with breakpoints off by at most 100 bp [and by at most 2,000 bp respectively]; and “N” indicates the method unable to detect the SV with breakpoints off by at most 2000 bp. SENSV can detect more SVs, especially for difficult cases, with much fewer false positives. Other software usually detects much more SVs than SENSV but most of them are false positives.

| ID | SV length | SV type | Detect the pathogenic SV? (# of predicted SVs of the same type) |  |  |  |
| --- | --- | --- | --- | --- | --- | --- |
|  |  |  | SENSV | NanoVar | Sniffles | SVIM |
| Difficult cases |  |  |  |  |  |  |
| 1 | 146K | long deletion | Y (1,513) | Y (1,278) | Y- (1,027) | Y (2,078) |
| 2 | 266K | long deletion | Y (1,431) | Y (1,353) | Y (1,172) | Y (2,129) |
| 3 | 638K | long deletion | N (1,743) | N (1,146) | N (732) | N (2,207) |
| 4 | 670K | long deletion | Y (1,612) | N (1,176) | N (794) | N (2,036) |
| 5 | 1.5M | long deletion | Y (3,395) | N (1,514) | N (1,062) | N (4,970) |
| 6 | 1.4M | long deletion | Y (17,209) | N (3,692) | N (1,375) | Y- (17,783) |
| 7 | 1.4M | duplication | Y (200) | N (1,779) | N (667) | Y- (1,901) |
| 8 | 2.8M | long deletion | Y (1,792) | N (1,320) | N (1,192) | N (2,983) |
| 9 | 5.2M | long deletion | Y (1,515) | N (1,294) | N (1,014) | N (1,982) |
| 10 | 6.6M | long deletion | Y (2,868) | N (1,474) | Y (1,362) | Y (4,534) |
| 11 | 342K | unbalanced translocation | N (490) | N (7,106) | N (1,383) | N (1,154) |
| 12 | 1.4M | unbalanced translocation | Y (462) | N (7,802) | N (1,432) | N (1,179) |
| 13 | 5.9M | terminal deletion | Y (3,476) | N (1,498) | N (1,510) | N (4,649) |
| 14 | 18M | unbalanced translocation | Y (269) | N (6,156) | N (1,719) | N (1,429) |
| 15 | 19M | terminal deletion | Y (1,413) | N (1,220) | N (968) | N (2,031) |
| 16 | 58M | unbalanced translocation | Y (883) | N (6,688) | N (2,340) | N (2,053) |
| Others |  |  |  |  |  |  |
| 17 | 142K | inversion | Y (114) | Y- (1,413) | Y- (1,391) | Y- (849) |
| 18 | 73M | inversion | Y (118) | Y (1,333) | Y (853) | Y (523) |
| 19 | 33M | inversion | Y (192) | Y (1,258) | N (190) | Y (117) |
| 20 | N/A | balanced translocation | Y (89) | Y (5,554) | Y (1,463) | Y (1,190) |
| 21 | N/A | balanced translocation | Y (861) | Y (6,898) | Y (2,594) | N (2,012) |
| 22 | N/A | balanced translocation | Y (76) | Y (5,254) | Y (1,187) | N (996) |
| 23 | N/A | balanced translocation | Y (111) | Y (5,542) | Y (1,449) | N (1,065) |
| 24 | N/A | balanced translocation | Y (592) | Y (6,628) | Y (1,775) | Y (1,358) |

Samples 11 to 16 concern an unbalanced translocation and terminal deletion that occurs near a highly repeated telomere region. They involve a deletion at highly repeated telomere regions on different chromosomes. They are difficult to detect because (1) a read might be aligned to the telomere region of different chromosomes with similar patterns, and (2) the genome reference of the telomere regions of some chromosomes are missing. SENSV can detect five out of six complicated unbalanced SVs (samples 13 to 16) by applying SV-DP to detect the correct alignment, filtering false negative alignment by depth information and reconstructing the genome reference of each chromosome’s telomere by assembling ONT reads in a 150x standard NA12878 sample. While the other software tools failed to detect these SVs.

Balanced SVs, which can be detected by a read covering one of the two breakpoints (unbalanced SV can be detected by a read covering exactly one breakpoint only), can usually be detected by existing software (sample 17 to 24). However, more false positive balanced SVs are predicted due to 1) chimeric reads introduced by sequencing error and misalignment, and 2) inability to filter false positives by sequencing depth information. Existing software usually predicts unexpectedly large number of translocations (hundreds to thousands) while a patient with genetic disease is expected to have fewer than 10 translocations^21^. This large number of likely false positive SVs will introduce difficulties of distinguishing the disease-causing balanced SVs from the false positives. SENSV, by applying realignment and filtering, limited the number of false positives within a few hundreds. It can reduce a large effort of distinguishing the disease-causing balanced SVs.

### Real Normal DNA Data

We also evaluated SENSV by sequencing the normal reference DNA sample HG002,. commonly used for evaluating SV calling software. Our evaluation is based on a recently published benchmark set^18^ containing 532 confirmed SV with size at least 1 Kbp. They, except one, are all short deletions with size shorter than 100 Kbp. The DNA sample was sequenced using a single MinION flow cell, generating 3.7x data. Table 2 shows the sensitivity of SENSV, NanoVar, Sniffles and SVIM on detecting SVs in HG002 with breakpoint precision of 100 bp and 2,000 bp respectively. For detecting short deletions with size smaller than 100 Kbp, the sensitivity of SENSV is slightly higher than SVIM and much better than NanoVar and Sniffles. As most of the confirmed SVs in HG002 are short deletions with an average length 4,259 bp, they can be detected easily (as the gap for a single split-read alignment is small). Notice that some software tools predicted 2 to 4 times more SVs than SENSV. As the SVs on HG002 are not all detected and confirmed by the society, we cannot draw any conclusion on the correctness of the predicted SVs. The number of SV reported by each software tool can be found in the Supplementary Material S.2.

**Table 2.** The performance of SENSV, NanoVar, Sniffles and SVIM on detecting SVs in HG002 with breakpoint precision of 100 bp and 2,000 bp respectively. The benchmark set of HG002 contains 531 confirmed short deletions with size of smaller than 100 Kbp and one long deletion with size of larger than 100 Kbp. The sensitivity inside the paratheses is measured with the relaxed breakpoint precision of 2,000 bp.

|  | Short deletions (<100 Kbp) |  | Long deletion (>100 Kbp) |  |
| --- | --- | --- | --- | --- |
|  | # of confirmed SV detected<br>±100bp (±2,000 bp) | Sensitivity<br>±100bp (±2,000 bp) | # of confirmed SV detected<br>±100bp (±2,000 bp) | Sensitivity<br>±100bp (±2,000 bp) |
| **SENSV** | **389 (431)** | **73.26% (81.17%)** | **1 (1)** | **100% (100%)** |
| **NanoVar** | 353 (381) | 66.48% (71.75%) | 0 (0) | 0% (0%) |
| **Sniffles** | 269 (313) | 50.66% (58.95%) | 0 (0) | 0% (0%) |
| **SVIM** | 378 (427) | 71.19% (80.41%) | **1 (1)** | **100% (100%)** |

### Simulated Patient Data

To further evaluate SENSV’s ability on detecting long deletions (>100 Kbp), duplications, terminal deletions and unbalanced translocations (i.e., the difficult cases) and evaluate the correctness of the predicted SVs, we generated two simulated “patient genomes”. Each “patient genome” was generated by implanting 35 heterozygous SVs on the human genome reference hg19. The 35 SVs consisted of 5 SV of the following 7 types: short deletion, long deletion, duplications, terminal deletions, inversions, balanced translocations and unbalanced translocations. All deletions and duplications were selected randomly from the clinically confirmed SVs in the dbVar^22^ database. As there are fewer than 10 confirmed terminal deletion, inversions, and balanced and unbalanced translocations in dbVar, these SV types were simulated using RSVSim^23^. The terminal deletions and unbalanced translocations were randomly selected from either the 5’ end or 3’ end of a chromosome with one of the breakpoints lies in the telomere region. All SVs were not overlapped. These SVs with known positions were implanted into the reference genome hg19 using RSVSim. NanoSim^24^ was used to generate 12 Gbp (i.e., depth of 4x) of WGS data for each “patient genome”. The maximum read length was set to 20 Kbp (similar to our library preparation protocol for real data). Other characteristics of reads, including mismatch rate and length distribution, were trained based on real public data: ONT WGS consortium rel6^25^.

We evaluated the performance of SENSV, NanoVar, Sniffles and SVIM on calling the implanted SVs (against the reference genome hg19) with the breakpoints error of 100 bp and 2,000 bp respectively. Table 3 shows the number of SV for each type could be detected by the software tools in the two simulated datasets. Overall speaking, SENSV detected more or equal numbers of implanted SVs for each type. For those SV types that are difficult to be detected, i.e., long deletion, duplication, terminal deletion and unbalanced translocation, SENSV outperformed other software significantly. With the advantages of utilization of the depth information, SENSV was able to recover some of the missing long deletions and duplications whose split-reads were not found by the aligners used by other software tools. As a result, it could detect 7 to 11 more deletions and duplications than other software tools. For terminal deletion and unbalanced translocation with one breakpoint lies in the telomere region, existing software tools usually cannot distinguish reads sequenced from different telomere regions of different chromosomes or different positions of in the telomere regions. SENSV could distinguish them based on a more accurate alignment of reads in the telomere region or the nearby sub-telomere region. Thus, SENSV can detect 14 (out of 20) terminal deletion and unbalanced translocation while the second best software tool, SVIM, could detect 4 only. For the other cases that can be detected relatively easier, i.e. short deletion, inversion and balanced translocation, SENSV shared similar performance with other software tools.

**Table 3.** The number of SVs detected with a breakpoint precision of 100 bp by the software grouped by SV types in two simulated datasets. The number in the parentheses is the number of SVs detected with a breakpoint precision of 2000 bp. Each type has total 10 SVs implanted in two simulated datasets.

|  | # of Detected SVs for the 10 implanted SVs |  |  |  |
| --- | --- | --- | --- | --- |
|  | SENS SV | NanoVar | Sniffles | SVIM |
| **Difficult cases** |  |  |  |  |
| Long deletion (>100 Kbp) | 9 | 6 | 3 (4) | 5 (6) |
| Duplication | 8 | 4 | 4 | 5 |
| Terminal deletion | 7 | 0 | 0 | 2 |
| Unbalanced translocation | 7 | 2 | 3 | 2 |
| **Others** |  |  |  |  |
| Short deletion (<100 Kbp) | 10 | 4 | 7 | 10 |
| Inversion | 9 | 9 | 9 | 5 |
| Balanced translocation | 8 | 8 | 7 | 3 |

We have also evaluated those SVs cannot be detected by SENSV, almost all of them were due to no reads being sequenced covering the breakpoints of the planted SVs. There is one exception that there is one split-read can cover the breakpoint of a balanced translocation missed by SENSV. Although SENSV can find the split-read alignment, it did not report the translocation because the alignment score is not good enough. The details of the performance for each SV in each simulated datasets can be found in the Supplementary Section S.1 and S.2.

## Method

SENSV is a computational method for detecting SVs using low-depth ONT WGS data. Figure 1 shows the workflow of SENSV, which has four major steps: (1) SENSV obtains SV candidates from the alignment information of raw reads to the human reference genome. (2) It compares the sequencing depth of each DNA region with 24 reference datasets to detect another set of SV candidates. (3) All these SV candidates are refined using SV-DP, an SV-aware dynamic programming algorithm implemented to find precise breakpoints of the SV. (4) The candidates are filtered and refined based on base-level alignment to a modified reference genome. Quality scores of called SVs are also available, making it possible to apply additional filtering.

**Figure 1.**
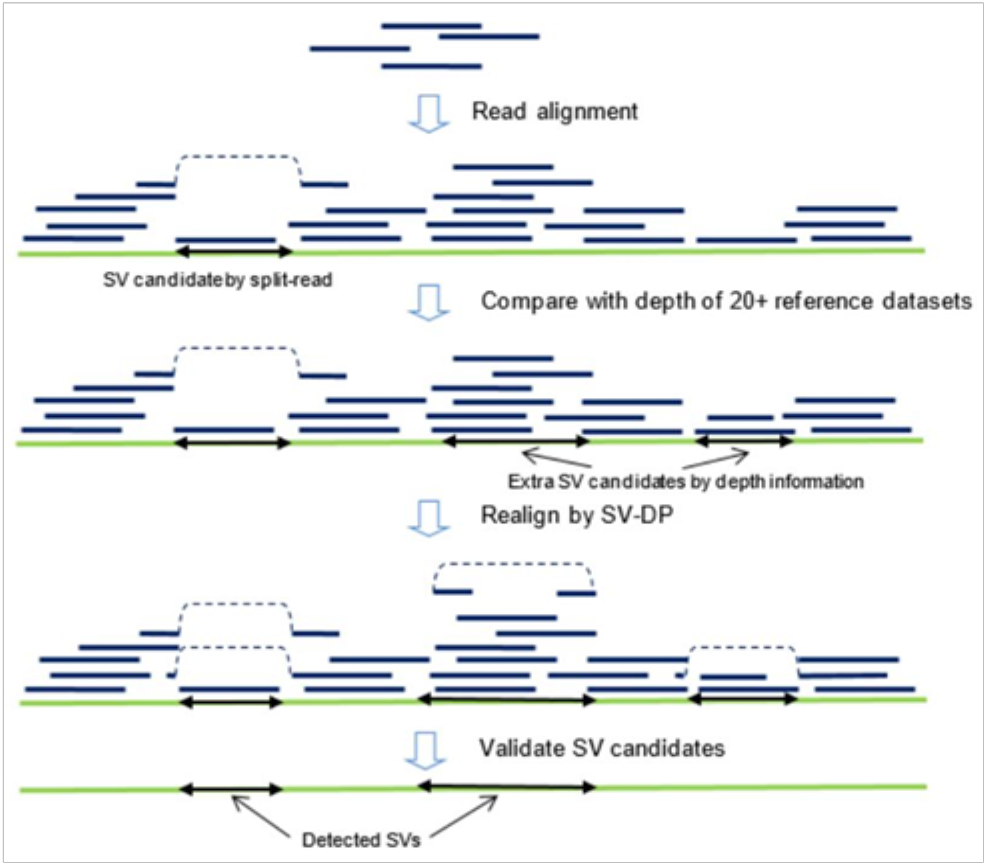
The workflow of SENSV.

SENSV makes use of Minimap2^19^ to align long reads to the human reference genome to find initial SV candidates. Minimap2 is very efficient in processing noisy long reads. A split-read alignment, which aligns the prefix and suffix of a read to different positions, is considered an SV candidate. Many SV candidates could be missed, as reads could be misaligned or poorly aligned, SENSV exploits other methods to recover and to verify the candidates. Note that existing software often cannot properly align a read near the end of a chromosome, as it might involve a highly repeated telomere region, and the DNA of such a region might be missing from the reference genome. Therefore, we used our own assembly of telomere sequence in SENSV. We aligned a 150x ONT WGS data of a standard human sample NA12878 from GIAB^26^ to the reference genome and found the reads aligned to para-telomere sequences of each chromosome. We assembled these reads using miniasm^27^ and used the assembled sequences as an additional reference DNA for read alignment in SENSV.

### SV Candidates via Sequencing Depth

Since fewer reads are sequenced in a deletion region and more reads in a duplication region, an unbalanced SV candidate, in principle, can be detected by analyzing the sequencing depth of each DNA region. However, when the sequencing depth is low (4x), the sequencing bias in different DNA regions may introduce many false positives and negatives. To solve this problem, SENSV considers only the DNA regions with abnormal depth compared with a reference dataset. The reference dataset is constructed by ONT WGS data from 24 people that shared no common SVs detected by Karyotyping and CGH array. The average depth of these 24 ONT WGS data is 4.3x (Supplementary Fig. S1) and the depth is normalized to 4x for each sample.

We partition the genome into intervals of 10 Kbp. The interval of 10 Kbp is chosen for the calculation as the average read length of the ONT WGS data is 10 Kbp. While we are working on low-depth samples, having an interval too short might result in insufficient or even no depth, while an interval too long might reduce breakpoint resolution. For each interval, the average depth is compared to that of the corresponding intervals of the 24 reference datasets. The comparison is accomplished by calculating the log-likelihood of two normal distribution models, representing the sequencing depth distribution of the sample with and without SV, respectively. We assume that each interval is independent of each other (in view of the interval size being sufficiently large), and define the two models as N(*μ*, *σ*^2^) and N(*μ*_SV_, *σ*^2^), where *μ* and *σ*^2^ represent the mean depth and standard deviation of the corresponding intervals in the 24 reference datasets, *μ*_SV_ is set to 0.5*μ* for detecting deletion and 1.5*μ* for detecting duplication (both assuming heterozygous SV). The likelihood ratio test is used to detect whether the depth of the analyzed interval is more likely to follow N(*μ*, *σ*^2^) or N(*μ*_SV_, *σ*^2^), i.e., to be normal or abnormal.

Lastly, nearby abnormal intervals are merged through a greedy algorithm to form a larger SV, and the rough breakpoints are generated. Since the likelihood ratio test follows the chi-square distribution, SENSV can calculate a *p*-value for the merged SV and consider regions with *p*-value ≤0.05 as SV candidates (in addition to the candidates detected by the split-read alignment). Note that this method can also detect homozygous SV because the distribution for an interval with a homozygous SV is closer to N(*μ*_SV_, *σ*^2^) than N(*μ*, *σ*^2^).

### SV-aware Aligner and Breakpoint Refinement

Many general-purpose aligners, such as Minimap2, calculate the alignment score based on a substitution matrix and gap-scoring scheme, and a long deletion is penalized by a large gap extension penalty. This penalty is not desirable in finding the breakpoints of SV. We propose an SV-aware aligner, called SV-DP, to find the breakpoints of large SVs.

SV-DP adopts a scoring scheme that does not penalize at most one gapped alignment introduced by an SV. By not penalizing the gapped alignment, which is caused by SV (including SVs other than deletion), the other parts of the read have a higher chance of being aligned correctly to the reference, thus obtaining more precise breakpoints.

The implementation of SV-DP is non-trivial because the time for aligning the read then depends on the size of the largest gap (i.e., the targeted one), which can be as big as several Mbp. To speed up SENSV (while retaining the same level of sensitivity), we perform SV-DP only on sequences near candidate breakpoints. Let *R* be a reference sequence, and let *Q* be a read sequence. Let *w* and *W* be predefined window sizes (2 Kbp and 10 Kbp, respectively). Assume *Q*[1..*q*] is aligned to *R*[*i*..*s*], and *Q*[*q*+1..*end*] to *R*[*e*..*j*] in the initial alignment of an SV candidate. We consider *Q*[*q*−*w* .. *q*+*w*] as a query and stitch two pieces of reference *R*[*s*−*w* .. *s*+*W*], *R*[*e*−*W*..*e*+*w*] together as the new reference. In this way, the running time of SV-DP is O(*wW*), which does not depend on SV size. We can recover the exact breakpoint, even if the candidate breakpoint is offset by *w* in the query and/or *W* in the reference.

For candidates found by sequencing depth, where the breakpoint is not accurate, SENSV reuses the seeding results from Minimap2 to find the supporting reads and narrows down the range of breakpoints. We extracted seeding information from Minimap2 in the format of [*query start*, *query end*, *ref start*, *ref end*]. For a candidate from depth analysis, if we can find one read with one seed near the starting breakpoint and another seed near the ending breakpoint, we proceed with this read to SV-DP to refine the breakpoint.

Finally, after all breakpoints are refined, de-duplication is applied to remove duplicate candidates.

### SV Validation

Once SENSV has figured out the breakpoints, the SV candidates can be validated using an alternative reference sequence. For a SV candidate, an alternative reference sequence is constructed by extracting nearby sequences from the reference genome and inserting the candidate into the sequence. If there are multiple SV candidates with similar breakpoints, multiple alternative reference sequences are constructed. If the SV candidate is genuine, we expect to see more reads previously aligned to the reference sequence aligned to the alternative sequence. Therefore, a candidate is discarded if its alternative reference fails to attract more reads. To minimize possible errors due to the existence of other SVs, the final set of SV candidates are validated again using an alternative reference sequence, as well as the entire human reference genome. If reads can be confidently aligned to the alternative reference rather than elsewhere in the human reference genome, it is likely that the corresponding candidate is genuine.

Finally, the QUAL score of the SV is derived from the alignment results in the validation process. Features like alignment length in query and reference and the number of matching bases are included for filtering the false positive SVs.

### Evaluation method

NanoVar was evaluated using the default parameters, with raw reads as input (NanoVar has an internal aligner). For Sniffles and SVIM, as recommended in their paper, we first performed the alignment using NGMLR^2^ and then used the generated bam file as inputs. Since the default parameters of the two software programs were not optimized for low-dept, to increase sensitivity, we consulted the developers of the software and tried multiple combinations of parameters. The best combination we found for the two software is shown in Supplementary Table S8. For SENSV, the initial alignment was done by Minimap2, and realignment using the SV-DP module was always performed.

SURVIVOR was primarily used to access the sensitivity of the software in detecting the SVs with the command “SURVIVOR eval” with the allowance of 100-bp or 2000-bp error distance from the breakpoints of the true-sets. This means only the predicted SVs with correct SV type prediction and the breakpoints being within 100-bp error are considered the true positives (TP). The number of SVs reported by each software is calculated by using “Bcftools filter”^28^ to only include the predicted SVs with certain sizes (greater than 1,000 bp) and SV types. In order to maximize the sensitivity for translocation, we also considered the predicted SVs which were classified as “BND”. The example commands used for SURVIVOR and bcftools can be found in the Supplementary Table S7.

## Conclusion

SENSV, by integrating several efficient algorithmic techniques, including SV-aware alignment (SV-DP), analysis of sequencing depth information, and sophisticated verification via re-alignment, can effectively utilize 4x ONT whole genome sequencing data to detect heterozygous structural variations with superior sensitivity, precision and breakpoint resolution. This makes clinical diagnosis of pathogenic SV using a single MinION flowcell feasible and cost effective.

## Data availability

The source code of SENSV is available on Github: github.com/HKU-BAL/SENSV, implemented in Python 3.

## Author contributions statement

H.L., H.Y. and Y.Z. conceived the presented ideas and experiments. H.Y. Y.Z and Y.W. conducted the benchmarking experiments. H.L. and H.Y. analyzed the results. W.L. performed the sequencing experiments. H.Y. Y.Z. K.M. and Y.W. wrote the code for SENSV. R.L. and T.L. supervised the project.

