## Supplementary Material for "SENSV: Detecting Structural Variations with Precise Breakpoints using Low-Depth WGS Data from a Single Oxford Nanopore MinION Flowcell"

### S.1. Performance of SENSV, NanoVar[1], Sniffles[2], and SVIM[3] on the simulated datasets for individual SV

| SV ID | SV length | SV type | SENSV | NanoVar | Sniffles | SVIM |
|---|---|---|---|---|---|---|
| 1 | 39,438 | deletion | Y | Y | Y | Y |
| 2 | 82,549 | deletion | Y | N | N | Y |
| 3 | 94,159 | deletion | Y | Y | Y | Y |
| 4 | 10,117 | deletion | Y | Y | Y | Y |
| 5 | 48,202 | deletion | Y | N | Y | Y |
| 6 | 33,546 | deletion | Y | N | N | Y |
| 7 | 23,876 | deletion | Y | N | N | Y |
| 8 | 97,991 | deletion | Y | Y | Y | Y |
| 9 | 34,940 | deletion | Y | N | Y | Y |
| 10 | 62,420 | deletion | Y | N | Y | Y |
| 11 | 113,720 | long deletion | Y | Y | Y | Y |
| 12 | 838,659 | long deletion | Y | Y | Y- | Y- |
| 13 | 498,160 | long deletion | Y | N | Y | N |
| 14 | 1,095,657 | long deletion | Y | N | N | Y |
| 15 | 412,281 | long deletion | Y | Y | N | N |
| 16 | 937,186 | long deletion | Y | Y | Y | Y |
| 17 | 509,473 | long deletion | Y | N | N | N |
| 18 | 1,063,340 | long deletion | N | N | N | N |
| 19 | 119,059 | long deletion | Y | Y | N | Y |
| 20 | 1,045,185 | long deletion | Y | Y | N | Y |
| 21 | 940,447 | inversion | Y | Y | Y | N |
| 22 | 930,177 | inversion | Y | Y | Y | Y |
| 23 | 985,020 | inversion | Y | Y | Y | Y |
| 24 | 43,362 | inversion | Y | Y | Y | Y |
| 25 | 43,233 | inversion | Y | Y | Y | Y |
| 26 | 40,819 | inversion | N | N | N | N |
| 27 | 45,234 | inversion | Y | Y | Y | N |
| 28 | 41,191 | inversion | Y | Y | Y | N |
| 29 | 909,789 | inversion | Y | Y | Y | Y |
| 30 | 940,949 | inversion | Y | Y | Y | N |
| 31 | 470,496 | duplication | Y | N | N | N |
| 32 | 80,807 | duplication | Y | Y | Y | Y |
| 33 | 116,822 | duplication | Y | N | Y | Y |
| 34 | 47,269 | duplication | Y | Y | Y | Y |
| 35 | 56,779 | duplication | Y | Y | N | N |

| SV ID | SV length | SV type | SENSV | NanoVar | Sniffles | SVIM |
|---|---|---|---|---|---|---|
| 36 | 896,748 | duplication | Y | N | N | Y |
| 37 | 1,189,698 | duplication | Y | N | Y | Y |
| 38 | 55,797 | duplication | N | N | N | N |
| 39 | 499,008 | duplication | N | N | N | N |
| 40 | 40,566 | duplication | Y | Y | N | N |
| 41 | 850,000 | terminal deletion | Y | N | N | N |
| 42 | 600,000 | terminal deletion | Y | N | N | N |
| 43 | 1,000,000 | terminal deletion | Y | N | N | N |
| 44 | 800,000 | terminal deletion | N | N | N | N |
| 45 | 900,000 | terminal deletion | Y | N | N | N |
| 46 | 2,500,000 | terminal deletion | N | N | N | N |
| 47 | 3,000,000 | terminal deletion | Y | N | N | Y |
| 48 | 1,500,000 | terminal deletion | Y | N | N | Y |
| 49 | 1,000,000 | terminal deletion | Y | N | N | N |
| 50 | 2,720,000 | terminal deletion | N | N | N | N |
| 51 | N/A | balanced translocation | Y | Y | Y | Y |
| 52 | N/A | balanced translocation | N | N | N | N |
| 53 | N/A | balanced translocation | Y | Y | Y | N |
| 54 | N/A | balanced translocation | Y | Y | Y | N |
| 55 | N/A | balanced translocation | N | Y | N | N |
| 56 | N/A | balanced translocation | Y | Y | Y | N |
| 57 | N/A | balanced translocation | Y | Y | Y | N |
| 58 | N/A | balanced translocation | Y | N | N | N |
| 59 | N/A | balanced translocation | Y | Y | Y | Y |
| 60 | N/A | balanced translocation | Y | Y | Y | Y |
| 61 | N/A | unbalanced translocation | Y | N | N | N |
| 62 | N/A | unbalanced translocation | Y | Y | Y | Y |
| 63 | N/A | unbalanced translocation | Y | N | N | N |
| 64 | N/A | unbalanced translocation | Y | N | N | N |
| 65 | N/A | unbalanced translocation | N | N | N | N |
| 66 | N/A | unbalanced translocation | Y | Y | Y | Y |
| 67 | N/A | unbalanced translocation | N | N | N | N |
| 68 | N/A | unbalanced translocation | Y | N | N | N |
| 69 | N/A | unbalanced translocation | N | N | N | N |
| 70 | N/A | unbalanced translocation | Y | N | Y | N |

**Table S1.** Comparison of SENSV, NanoVar, Sniffles and SVIM on the ability to detect the SV in the simulated datasets. Above, “Y” [and “Y-”] mean that a software can detect the SV with the correct SV type and with a breakpoint precision of 100 bp [and of 2,000 bp]; and “N” indicates the software is unable to detect the SV with breakpoint precision relaxed to 2,000 bp.

### S.2. The number of reported SV in different types detected by the software for the datasets

Deleted: S.1. Performance of SENSV, NanoVar, Sniffles, and SVIM on the simulated datasets with different depth¶
¶
¶
... [1]

| | Number of predicted deletions | | | |
|---|---|---|---|---|
| Dataset ID | SENSV | NanoVar | Sniffles | SVIM |
| HG002 | 1,850 | 1,209 | 984 | 2,280 |
| simulated data A | 81 | 37 | 52 | 200 |
| simulated data B | 55 | 26 | 46 | 212 |
| 1 | 1,513 | 1,278 | 1,027 | 2,078 |
| 2 | 1,431 | 1,353 | 1,172 | 2,129 |
| 3 | 1,743 | 1,146 | 732 | 2,207 |
| 4 | 1,612 | 1,176 | 794 | 2,036 |
| 5 | 3,395 | 1,514 | 1,062 | 4,970 |
| 6 | 17,209 | 3,692 | 1,375 | 17,783 |
| 7 | 1,353 | 1,192 | 973 | 1,715 |
| 8 | 1,792 | 1,320 | 1,192 | 2,983 |
| 9 | 1,515 | 1,294 | 1,014 | 1,982 |
| 10 | 2,868 | 1,474 | 1,362 | 4,534 |
| 11 | 1,675 | 1,282 | 805 | 2,489 |
| 12 | 2,195 | 1,428 | 779 | 2,832 |
| 13 | 3,476 | 1,498 | 1,510 | 4,649 |
| 14 | 35,848 | 4,137 | 2,118 | 35,169 |
| 15 | 1,413 | 1,220 | 968 | 2,031 |
| 16 | 2,275 | 1,355 | 1,301 | 3,683 |
| 17 | 3,930 | 1,437 | 1,657 | 6,666 |
| 18 | 1,750 | 1,404 | 1,294 | 2,781 |
| 19 | 1,100 | 1,165 | 455 | 973 |
| 20 | 1,543 | 1,211 | 964 | 1,767 |
| 21 | 2,197 | 1,423 | 1,359 | 3,411 |
| 22 | 1,195 | 1,183 | 786 | 1,425 |
| 23 | 2,002 | 1,317 | 965 | 2,085 |
| 24 | 2,218 | 1,366 | 992 | 2,670 |

**Table S2.** The number of reported deletions by each software.

| | Number of predicted duplications | | | |
|---|---|---|---|---|
| Dataset ID | SENSV | NanoVar | Sniffles | SVIM |
| HG002 | 285 | 1,882 | 759 | 2,730 |
| simulated data A | 31 | 27 | 36 | 216 |
| simulated data B | 29 | 40 | 41 | 242 |
| 1 | 190 | 2,225 | 784 | 2,191 |
| 2 | 209 | 2,502 | 903 | 2,370 |
| 3 | 140 | 1,568 | 514 | 2,006 |
| 4 | 190 | 1,837 | 572 | 1,950 |
| 5 | 276 | 3,017 | 776 | 3,501 |
| 6 | 218 | 2,962 | 755 | 2,042 |
| 7 | 200 | 1,779 | 667 | 1,901 |
| 8 | 200 | 2,236 | 933 | 3,120 |
| 9 | 160 | 2,562 | 697 | 1,914 |
| 10 | 276 | 3,030 | 1,085 | 3,805 |
| 11 | 139 | 2,128 | 599 | 2,075 |
| 12 | 137 | 2,245 | 621 | 2,195 |
| 13 | 333 | 4,177 | 1,246 | 3,825 |
| 14 | 283 | 2,769 | 843 | 2,692 |
| 15 | 154 | 2,208 | 704 | 2,064 |
| 16 | 236 | 2,606 | 972 | 3,591 |
| 17 | 506 | 3,672 | 1,145 | 6,787 |
| 18 | 182 | 2,662 | 872 | 3,159 |
| 19 | 149 | 1,610 | 247 | 928 |
| 20 | 175 | 2,062 | 610 | 1,765 |
| 21 | 246 | 2,670 | 935 | 4,037 |
| 22 | 155 | 1,756 | 535 | 1,556 |
| 23 | 157 | 2,244 | 665 | 1,838 |
| 24 | 240 | 2,572 | 679 | 2,240 |

**Table S3.** The number of reported duplications by each software.

| | Number of predicted inversions | | | |
|---|---|---|---|---|
| Dataset ID | SENSV | NanoVar | Sniffles | SVIM |
| HG002 | 233 | 1,261 | 634 | 351 |
| simulated data A | 11 | 20 | 69 | 115 |
| simulated data B | 7 | 24 | 85 | 129 |
| 1 | 141 | 1,194 | 600 | 439 |
| 2 | 72 | 1,360 | 708 | 463 |
| 3 | 103 | 975 | 478 | 428 |
| 4 | 159 | 1,076 | 448 | 394 |
| 5 | 167 | 1,280 | 700 | 502 |
| 6 | 256 | 1,256 | 594 | 420 |
| 7 | 96 | 1,071 | 573 | 329 |
| 8 | 226 | 1,194 | 776 | 574 |
| 9 | 176 | 1,240 | 597 | 404 |
| 10 | 411 | 1,381 | 861 | 645 |
| 11 | 219 | 1,153 | 444 | 324 |
| 12 | 125 | 1,236 | 476 | 370 |
| 13 | 271 | 1,615 | 986 | 697 |
| 14 | 233 | 1,244 | 724 | 579 |
| 15 | 240 | 1,163 | 636 | 453 |
| 16 | 167 | 1,286 | 897 | 660 |
| 17 | 114 | 1,413 | 1,391 | 849 |
| 18 | 118 | 1,333 | 853 | 523 |
| 19 | 192 | 1,258 | 190 | 117 |
| 20 | 207 | 1,112 | 566 | 346 |
| 21 | 160 | 1,437 | 1,026 | 633 |
| 22 | 98 | 1,052 | 444 | 255 |
| 23 | 177 | 1,275 | 545 | 303 |
| 24 | 237 | 1,278 | 589 | 329 |

**Table S4.** The number of reported inversions by each software.

| | Number of predicted translocations (reported as "BND") | | | |
|---|---|---|---|---|
| Dataset ID | SENSV | NanoVar | Sniffles | SVIM |
| HG002 | 1,276 | 8,422 | 1,609 | 1,289 |
| simulated data A | 21 | 150 | 61 | 14 |
| simulated data B | 20 | 170 | 65 | 12 |
| 1 | 116 | 5,720 | 1,543 | 1,363 |
| 2 | 101 | 6,238 | 1,718 | 1,581 |
| 3 | 209 | 5,942 | 1,185 | 941 |
| 4 | 91 | 5,254 | 1,206 | 1,110 |
| 5 | 1,101 | 8,832 | 2,035 | 1,690 |
| 6 | 164 | 5,938 | 1,547 | 1,284 |
| 7 | 92 | 5,230 | 1,375 | 1,076 |
| 8 | 867 | 6,998 | 2,138 | 1,917 |
| 9 | 80 | 5,672 | 1,501 | 1,334 |
| 10 | 751 | 6,970 | 2,265 | 1,964 |
| 11 | 490 | 7,106 | 1,383 | 1,154 |
| 12 | 462 | 7,802 | 1,432 | 1,179 |
| 13 | 682 | 7,058 | 2,486 | 2,085 |
| 14 | 269 | 6,156 | 1,719 | 1,429 |
| 15 | 128 | 5,512 | 1,401 | 1,182 |
| 16 | 883 | 6,688 | 2,340 | 2,052 |
| 17 | 972 | 7,378 | 3,358 | 2,505 |
| 18 | 1,053 | 6,976 | 2,451 | 1,826 |
| 19 | 87 | 4,904 | 655 | 382 |
| 20 | 89 | 5,554 | 1,463 | 1,190 |
| 21 | 861 | 6,898 | 2,594 | 2,012 |
| 22 | 76 | 5,254 | 1,187 | 996 |
| 23 | 111 | 5,542 | 1,449 | 1,065 |
| 24 | 592 | 6,628 | 1,775 | 1,358 |

**Table S5.** The number of reported translocations by each software.

| | Total | | | |
|---|---|---|---|---|
| Dataset ID | SENSV | NanoVar | Sniffles | SVIM |
| HG002 | 3,644 | 12,774 | 3,986 | 6,650 |
| simulated data A | 144 | 234 | 218 | 545 |
| simulated data B | 111 | 260 | 237 | 595 |
| 1 | 1,960 | 10,417 | 3,954 | 6,071 |
| 2 | 1,813 | 11,453 | 4,501 | 6,543 |
| 3 | 2,195 | 9,631 | 2,909 | 5,582 |
| 4 | 2,052 | 9,343 | 3,020 | 5,490 |
| 5 | 4,939 | 14,643 | 4,573 | 10,663 |
| 6 | 17,847 | 13,848 | 4,271 | 21,529 |
| 7 | 1,741 | 9,272 | 3,588 | 5,021 |
| 8 | 3,085 | 11,748 | 5,039 | 8,594 |
| 9 | 1,931 | 10,768 | 3,809 | 5,634 |
| 10 | 4,306 | 12,855 | 5,573 | 10,948 |
| 11 | 2,523 | 11,669 | 3,231 | 6,042 |
| 12 | 2,919 | 12,711 | 3,308 | 6,576 |
| 13 | 4,762 | 14,348 | 6,228 | 11,256 |
| 14 | 36,633 | 14,306 | 5,404 | 39,869 |
| 15 | 1,935 | 10,103 | 3,709 | 5,730 |
| 16 | 3,561 | 11,935 | 5,510 | 9,986 |
| 17 | 5,522 | 13,900 | 7,551 | 16,807 |
| 18 | 3,103 | 12,375 | 5,470 | 8,289 |
| 19 | 1,528 | 8,937 | 1,547 | 2,400 |
| 20 | 2,014 | 9,939 | 3,603 | 5,068 |
| 21 | 3,464 | 12,428 | 5,914 | 10,093 |
| 22 | 1,524 | 9,245 | 2,952 | 4,232 |
| 23 | 2,447 | 10,378 | 3,624 | 5,291 |
| 24 | 3,287 | 11,844 | 4,035 | 6,597 |

**Table S6.** The number of total reported SV by each software.

### S.3. The Algorithm of SV-DP

Let Q = q1q2 ... qn and R = r1r2...rn be query sequence and reference sequence respectively, where n and m are the lengths of the sequences.

Substitution matrix s(q,r) defines the similarity score of two nucleotides q and r,
and penalty function W(k) defines the penalty of a gap (deletion or insertion) with length k.

Scoring matrix H of our dynamic programming is a 3 dimensional table, and consists of 4 subtables.

H[k][i][j] represents the best alignment of Q[1..i] and R[1..j], with exactly k of the deletions has a zero penalty. In our SV detection application, only 1 free deletion is needed, and k will be either 0 or 1. In implementation, we use a circular buffer of size 2 (instead of size = query length) to represent the second dimension.

H_M[k][i][j] stores the best alignment that ends in match/mismatch (i.e. Q[i] is aligned to R[j])

H_I[k][i][j] stores the best alignment that ends in insertion (i.e. Q[i] is in an insertion), while insertion length is recorded in a separate table I_len[k][j][i].

H_D[k][i][j] stores the best alignment that ends in deletion (i.e. R[j] is in a deletion), while deletion length is recorded in a separate table D_len[k][j][k]

H_F[k][i][j] stores the best alignment that ends in deletion and the deletion is of free penalty.

For all these 4 subtables, we stores not only scores, but also the positions of the deletion with free penalty, if any (starting position in reference, ending position in reference, query position).

We keep track of the best alignment while filling the scoring matrices and therefore a backtracking would not be necessary to retrieve the deletion with free penalty.

Initialization:

```
H_M[k][j][i] = 0,
H_I[k][j][i] = -inf,
H_D[k][j][i] = -inf,
```

```
H_F[k][j][i] = -inf,

I_len[k][j][i] = D_len[k][j][i] = -inf

Transitions:

for i>1, j>1

(H_M is written as M)

I[k][i][j] = max(M[k][i-1][j] + W(1), I[k][i-1][j] + W(l+1) -
W(l)) where l = I_len[k][i-1][j]
D[k][i][j] = max(M[k][i][j-1] + W(1), D[k][i][j-1] + W(l+1) -
W(l)) where l = D_len[k][i][j-1]
if k>0:
    F[k][i][j] = max(M[k-1][i][j-1], F[k-1][i][j-1])
M[k][i][j] = max(I[k][i][j], D[k][i][j], F[k][i][j], M[k][i-
1][j-1] + s(Q[i],R[j]))

(supplementary information like I_len, D_len, and position of
deletion of free penalty are updated accordingly.)

RET = max(RET, M[k][i][j])

Return Value:
RET
```

### S.4. Depth variability of the reference dataset in SENSV

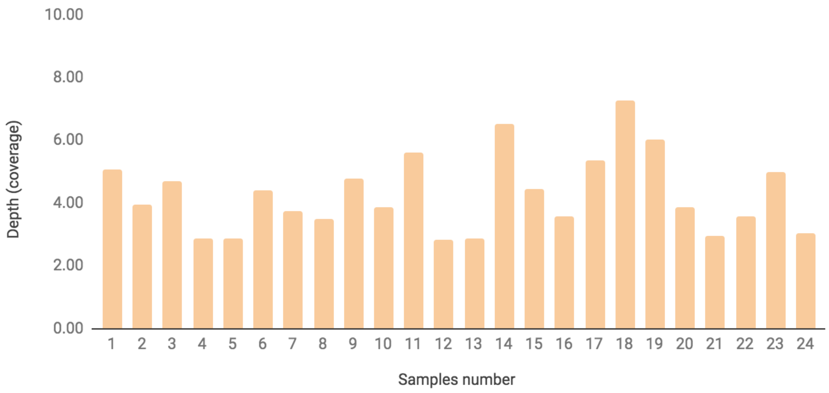

**Figure S1.** Sequencing depth of the samples for the reference dataset in SENSV

### S.5. Commands and parameters used for the evaluation

Deleted: Section Break (Next Page)

| Software | Command |
| --- | --- |
| SURVIVOR[4] (1.0.7) | SURVIVOR eval --vcf_results --true_set_bed2_format --100 (allowed error at the breakpoints) --output_dir |
| bcftools[5] | bcftools filter --i ‘SVTYPE==(Type)’--vcf \| bcftools filter --i ‘(END-POS)>=1000’ |

**Table S7**. Commands and parameters used for counting the number of reported SV and true positives.

### S.6.Parameters and software version used for calling SV for low-depth (4x) WGS data

| Software | Parameters |
|---|---|
| SENSV | --min_sv_size 1000 |
| Sniffles (1.0.11) | -s 2 |
| SVIM (1.1.1) | alignment --max_sv_size 250000000 |
| NanoVar (1.2.6) | -r hs37d5.fa |

**Table S8.** Parameters and versions used for the software when evaluating low-depth WGS data.

Formatted Table

**Page 5: [1] Deleted** **SR** **12/16/20 4:40:00 PM**
